# Miniaturized Devices for Bioluminescence Imaging in Freely Behaving Animals

**DOI:** 10.1101/2020.06.15.152546

**Authors:** Dmitrijs Celinskis, Nina Friedman, Mikhail Koksharov, Jeremy Murphy, Manuel Gomez-Ramirez, David Borton, Nathan Shaner, Ute Hochgeschwender, Diane Lipscombe, Christopher Moore

## Abstract

Fluorescence miniature microscopy *in vivo* has recently proven a major advance, enabling cellular imaging in freely behaving animals. However, fluorescence imaging suffers from autofluorescence, phototoxicity, photobleaching and non-homogeneous illumination artifacts. These factors limit the quality and time course of data collection. Bioluminescence provides an alternative kind of activity-dependent light indicator. Bioluminescent calcium indicators do not require light input, instead generating photons through chemiluminescence. As such, limitations inherent to the requirement for light presentation are eliminated. Further, bioluminescent indicators also do not require excitation light optics: the removal of this component should make lighter and lower cost microscope with fewer assembly parts. While there has been significant recent progress in making brighter and faster bioluminescence indicators, parallel advances in imaging hardware have not yet been realized. A hardware challenge is that despite potentially higher signal-to-noise of bioluminescence, the signal strength is lower than that of fluorescence. An open question we address in this report is whether fluorescent miniature microscopes can be rendered sensitive enough to detect bioluminescence. We demonstrate this possibility *in vitro* and *in vivo* by implementing optimizations of the UCLA fluorescent miniscope. These optimizations yielded a miniscope (BLmini) which is 22% lighter in weight, has 45% fewer components, is up to 58% less expensive, offers up to 15 times stronger signal (as dichroic filtering is not required) and is sensitive enough to capture spatiotemporal dynamics of bioluminescence in the brain with a signal-to-noise ratio of 34 dB.

## I. Introduction

Microscopic imaging of fluorescent indicators to track neural activity is an increasingly important tool in systems neuroscience. Over the last 9 years, portable microscopes that allow free behavior while collecting 1-photon signals have become regularly employed in freely behaving animals. This key advance became possible with the introduction of miniature microscopy (or ‘miniscopes’) in 2011 by Schnitzer and colleagues [1]. The adoption of this technology was accelerated around 2016 when a group from UCLA released their first version of open-source miniature microscope most commonly known as UCLA miniscope. [2] Ever since its release, UCLA miniscope has been used by over 400 different groups to answer questions in domains ranging from memory representation to olfactory processing. [3]

Despite their impact, fluorescence-based imaging tools are fundamentally limited by their need to project light onto the brain. A common problem that limits imaging sessions is photobleaching. A parallel common problem that limits the long-term survival of imaging fields is phototoxicity. The signal-to-noise ratio for localizing discrete signals in the tissue is also fundamentally limited, due to autofluorescence from unintended sources, and non-homogeneous illumination due to the scattering of excitation light. These issues become further exacerbated in the case of miniscopes, due to smaller form and cost factors compared to traditional benchtop microscopes. The weight requirement of mobile devices implanted in mice [4] inevitably constrains the optics, imaging quality and experimental duration attainable with epifluorescence miniscopes.

Bioluminescent molecular tools offer a promising alternative to fluorescent indicators. Bioluminescence is a form of “cold-light” generated via chemiluminescent reaction between a luciferase (an enzyme) and luciferin (a substrate). Many forms of bioluminescence in nature require a second factor to produce photons, such as increased ATP levels [5] or calcium concentration, [6] and there has been a recent increase in the synthesis of optimized luciferases for calcium sensing. [7]

Bioluminescent indicators create light within the cell of interest, and do not require excitation light, creating several benefits. Bioluminescent cells do not suffer from photobleaching, phototoxicity, or excitation scattering, and these point sources do not create autofluorescence. As such, the bioluminescent indicator strategy significantly reduces the noise ‘denominator’ of the SNR underlying definitive localization. Also, because excitation light is not needed, the form of the miniscope can, in concept, be made significantly simpler and lower weight by removing components associated with excitation. These gains are important when considering the challenge of *in vivo* mouse imaging.

Natural bioluminescent indicators were among the first employed in neuroscience [8]. However, native bioluminescent sources had two key limitations. First, many photoproteins required achieving a complex molecular configuration, causing fast rundown of the signal. These molecules also often had slow time constants and lower total signal output. These issues have largely been addressed by recent synthetic bioluminescent indicators, but whether their signal strength is sufficient for large-scale neural imaging *in vivo* remains a major question.

To directly address this key question, we systematically modified miniscope design for bioluminescence, and tested its capacity for detecting bioluminescent signals *in vitro* and *in vivo*. We found that this modified miniscope could robustly detect these signals, and when compared to a conventional system showed a significant increase in signal detectability, reduction in complexity, and decrease in weight. This progress represents a key step in the refinement of this approach, which can prove transformative to an important imaging approach.

## II. Methods

### A. Design of Bioluminescence Miniscope (BLmini)

The chemiluminescent generation of bioluminescence removes the need for excitation light optics, allowing beneficial simplification of microscope’s architecture. In the case of UCLA miniscope v3.2, we removed the excitation LED PCB, filters, dichroic mirror and filter set cover (Fig. 1).

**Figure 1.**
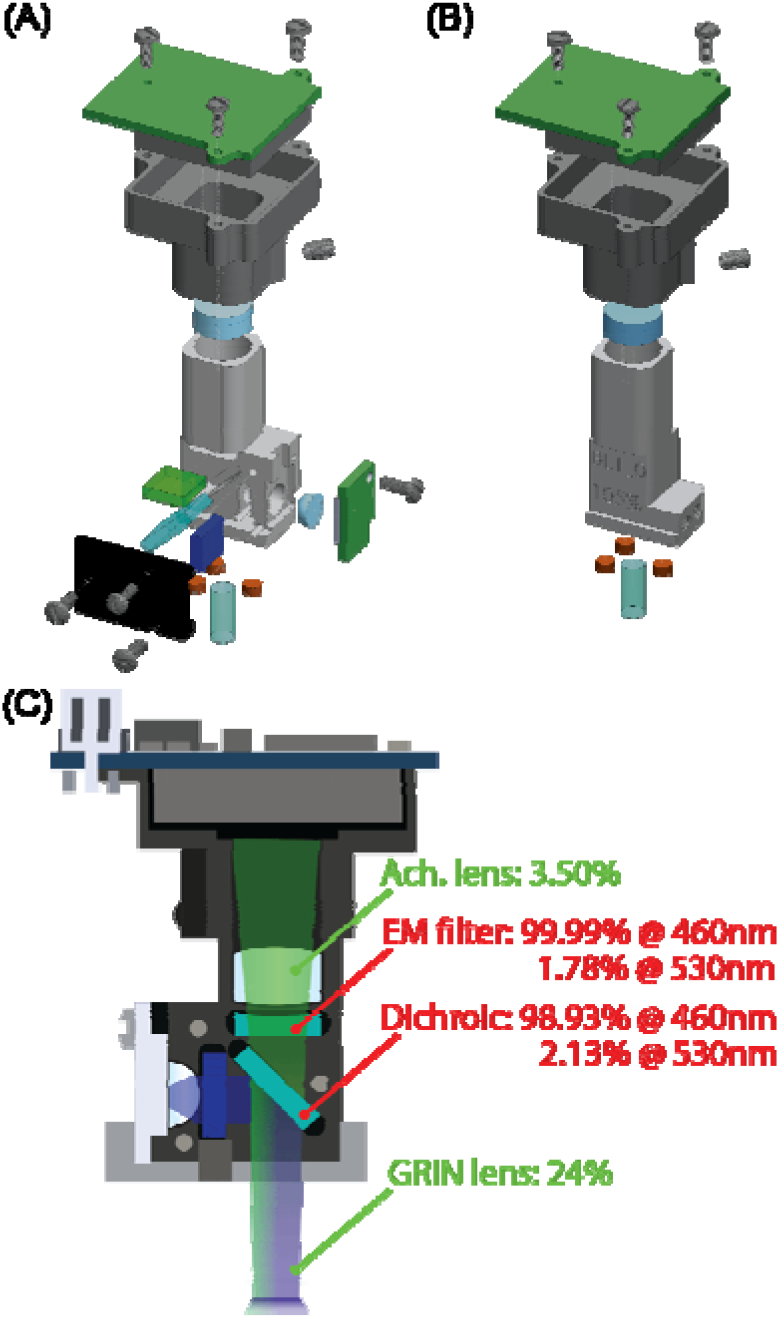
(A) An epifluorescence UCLA miniscope v3.2 [9] and (B) the optimized bioluminescence miniscope (BLmini) v1.0. (C) Cross-section of UCLA miniscope with percentage total light losses caused by the optical components along emission light path. The components marked in red were removed as part of design optimization for bioluminescence imaging.

### B. Arrangement for the In Vitro comparison of the standard UCLA system and the BLmini

The ability to detect light of the same intensity was compared between the two miniscopes as shown in Fig. 2. Both miniscopes were placed into a Petri dish with RIN lenses (#64-519, Edmund Optics) dipped into a luciferase solution containing 150 uL of either Nanoluc (NLuc, also referred as NanOgluc; synthesized by Twist Bioscience as a part of 10,000 Free Genes Project) or mVenus2-Nluc, corresponding to 460 nm and 530 nm peak emission wavelengths, respectively. To initiate bioluminescence emission, ~150 uL of luciferin (bisCTZ) diluted to 10-15 uM was added. All measurements were done inside a custom-made light-tight chamber. An optical powermeter (PM100D with S130VC sensor, ThorLabs) was used to validate bioluminescence emission and control for background light emission in the dark enclosure.

**Figure 2.**
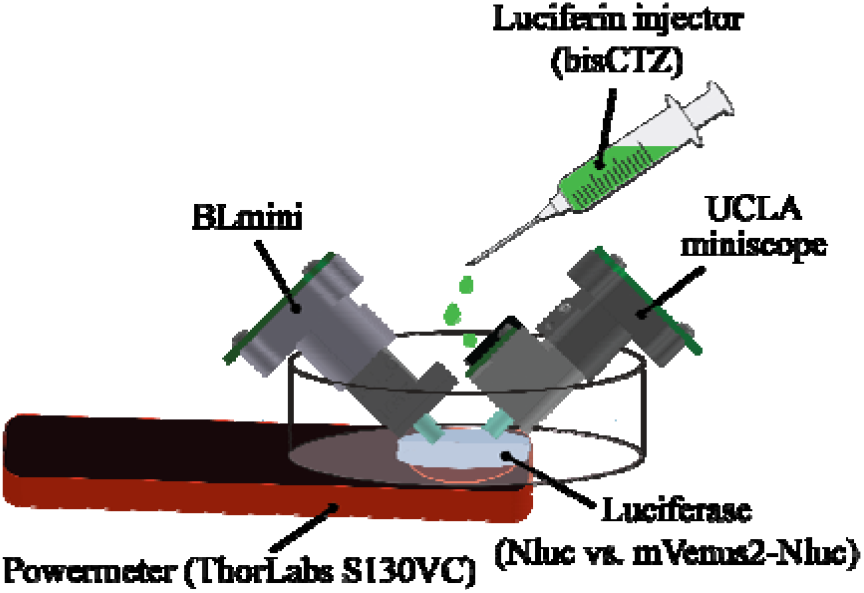
Experimental setup used for *in vitro* comparison of UCLA miniscope and BLmini. Bioluminescence signal was simultaneously tr cked using a powermeter underneath a Petri dish and two miniscopes immersed in the luciferase solution.

Miniscope and powermeter signals were synchronized in time using LED flashes. By default, the UCLA miniscope excitation LED is constantly turned on, even when set to lowest power. To avoid confounding light contamination, the excitation LED was disconnected from the sensor PCB during these measurements. Using UCLA miniscope software, both miniscopes were configured to sample at 5 fps (exposure time ~200 ms) and the gain set to its maximum of 64, a gain of 4 of the CMOS sensor (MT9V032C12STM, ON Semiconductor). To compute light intensities using acquired miniscope movies, the pixel intensity was averaged across all the pixels and normalized by the baseline pixel intensity.

### C. In Vivo Comparison of the UCLA standard configuration and the BLmini

*In vivo* testing was done on signals emanating from a mouse expressing mNeonGreen tethered to EkL9H luciferase (i.e. a shrimp-based luciferase variant also referred to as NCS2) 11 weeks following the injection of pAAV-hSyn-Kozak-NCS2 virus in primary somatosensory cortex (SI) (craniotomy centered at A/P = −1.25 mm and M/L = 3.25 mm relative to Bregma, injection depth = 0.35 mm). Before the experiment, dura was removed around the imaging area and a delivery pipette (34G, #207434, HAMILTON) brought into contact with the surface of the cortex, causing it to dimple. The injection of 1 uL of h-CTZ (2.36 mM, Cat # 3011, NanoLight Technologies) was carried out at a rate of 1.25 uL/min. A GRIN lens (GT-IFRL-180_inf_50-NC) mounted on the BLmini was lowered to touch the surface of the brain near the pipette. Using ambient light, we acquired brightfield images of the cortical surface to verify the location of the GRIN lens within the craniotomy based on vascular landmarks.

The BLmini was configured for image acquisition at 1 fps and software gain was set to its maximum value of 64. The miniscope was powered 40 min prior to imaging onset to minimize the impact of thermal noise. All measurements were conducted inside a custom-made dark enclosure with an animal under isoflurane anesthesia. An EMCCD camera (Ixon 888, Andor) with a Navitar Zoom 6000 lens system (Navitar, 0.5x lens) was used to acquire higher amplification cortical maps before and after imaging with the BLmini. The exposure time of the EMCCD camera was set to 10 s, and the EM gain was set to 30. Images from the BLmini and EMCCD camera were aligned manually based on vascular landmarks and used to verify the correspondence between peak signals picked up by two cameras.

## III. Results

### A. Greater signal strength, reduced mass, cost, power consumption and assembly complexity of the BLmini

Table 1 summarizes characteristics of the two miniscopes. The signal strength comparison was based on the experiment summarized in Fig. 2 and results are shown in Fig. 3. Both miniscopes were able to detect bioluminescence from both constructs, but the BLmini showed significantly stronger signal. The power budget in Table 1 was estimated for a wire-free version of UCLA miniscope released on 5/21/2019. [9]

**TABLE I.** Summary of Comparison of the Conventional UCLA miniscope and the BLmini

|  | UCLA miniscope | BLmini |
| --- | --- | --- |
| Mass | 3.2 g | 2.5 g |
| Number of components | 22 | 12 |
| Cost/scope | \$1263 (1-4 scopes)<br>\$547 ($\geq 20$ scopes) | \$526 (1-4 scopes)<br>\$443 ($\geq 20$ scopes) |
| Signal strength | x @ 460nm<br>y @ 530nm | $\sim 15x$ @ 460nm<br>$\sim 3y$ @ 530nm |
| Power consumption | 1,100 mW (wired)<br>296 mW (wire-free) | 957 mW (wired)<br>148 mW (wire-free) |

**Figure 3.**
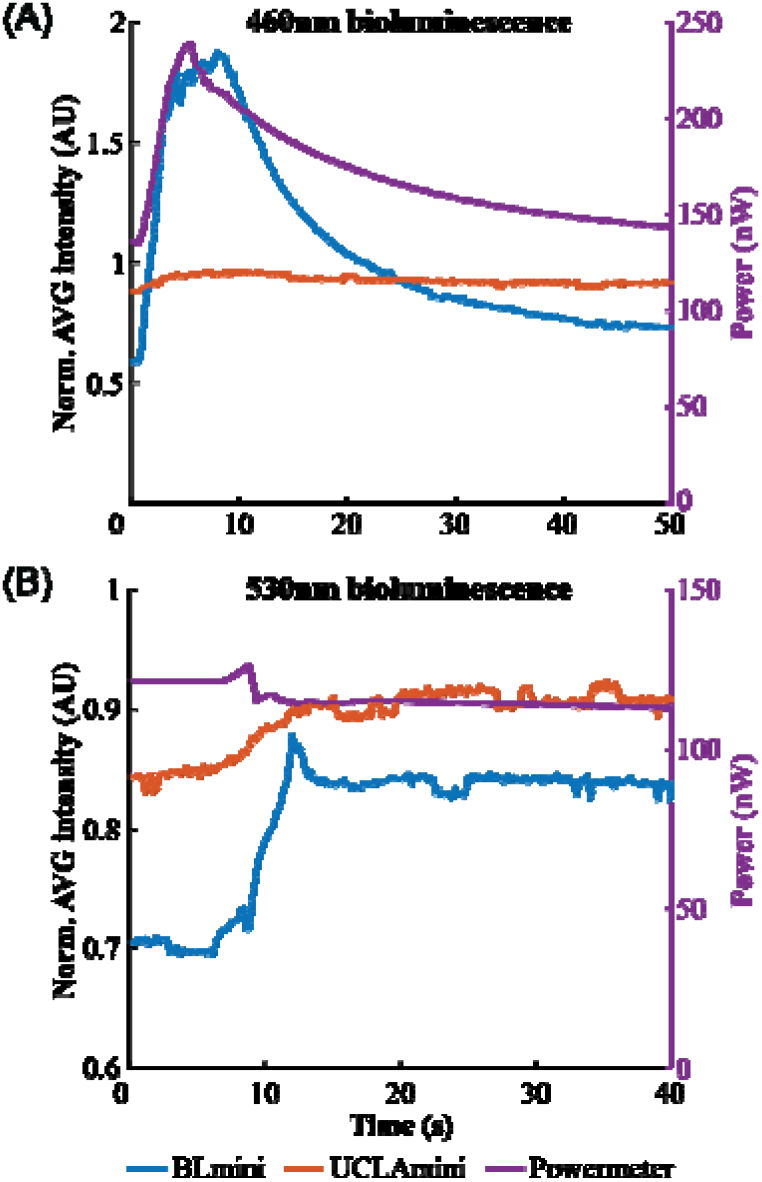
Results of *in vitro* bioluminescence measurements using the BLmini (blue), fluorescence miniscope (orange) and powermeter (purple) for two luciferase constructs. Besides differences in wavelengths ((A) for 460nm from Nluc and (B) for 530nm peak from mVenus2-Nluc), t o different luciferases had different emission intensities.

### B. Initial In Vivo Measurements with the BLmini Capture Temporal Dynamics of Bioluminescence

The BLmini was also able to capture the temporal dynamics of *in vivo* bioluminescence, as shown in Fig. 4. For comparison, we overlaid time courses acquired using the BLmini and by EMCCD in a different experiment ([10]; Fig. 4A). The signal captured by the BLmini also tracked more subtle features of the known BL response. These properties include an early slowing in photon production during substrate injection, thought to be due to inhibition of photon production by the breakdown products.

**Figure 4.**
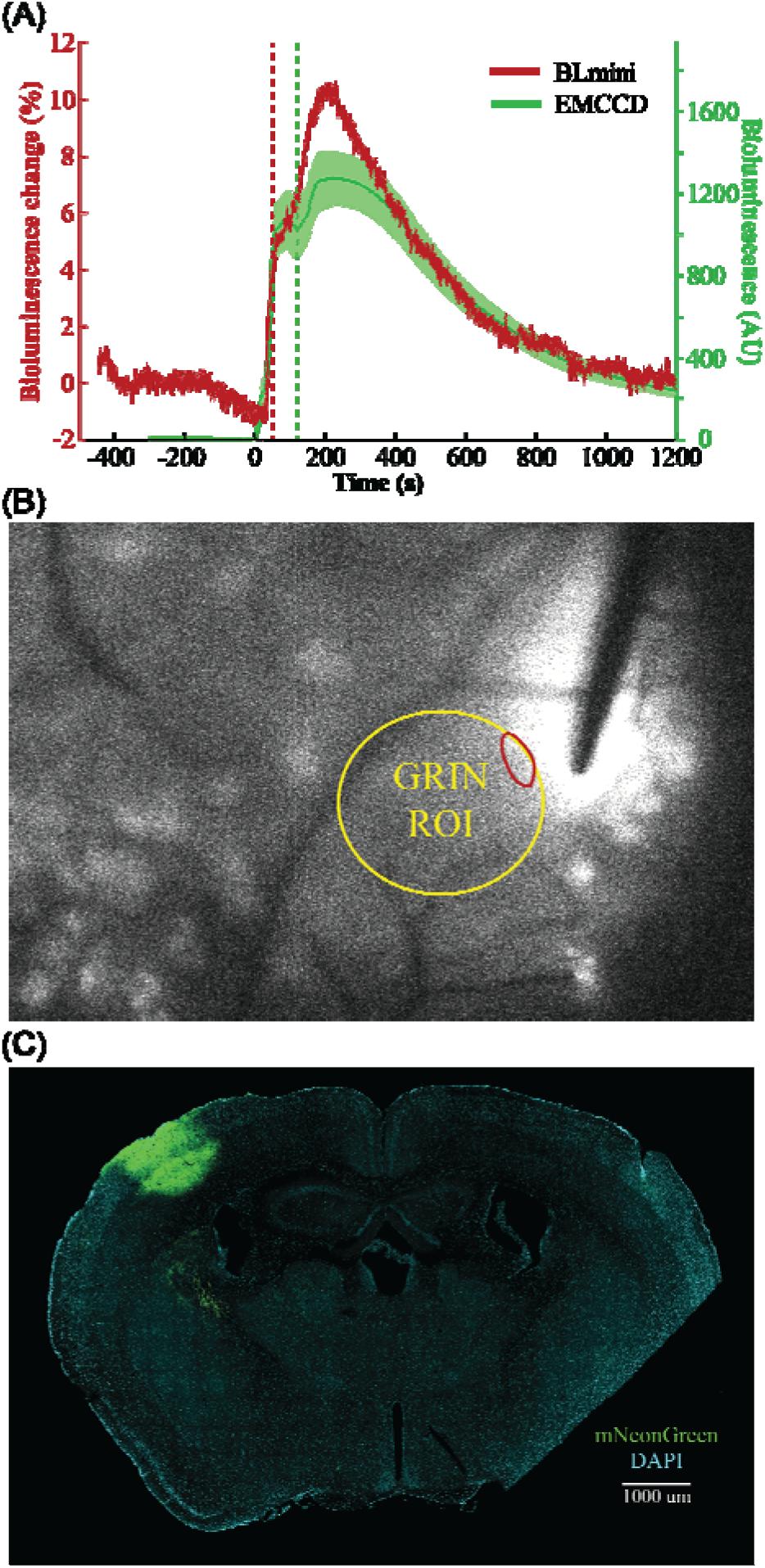
(A) Time courses of bioluminescence emission measured using BLmini (red) and EMCCD camera (green) in two different animals. BLmini imaging involved topical administration of h-CTZ and EMCCD imaging involved cortical injection of CTZ. (B) An image showing a bioluminescence map acquired using EMCCD camera immediately following BLmini measurements after removing the miniscope and GRIN lens from the field of view. Yellow ROI corresponds to an approximate prior location of GRIN lens and smaller red region corresponds to an ROI for which the mean intensity was plotted in (A). (C) Confocal microscopy image showing viral expression of mNeonGreen tethered to EkL9H luciferase on the coronal brain slice.

A clear difference between the EMCCD and miniscope lies in the baseline signal magnitude. CMOS sensor employed in the miniscope does not rely on any form of cooling and consequently suffers from significantly stronger thermoelectric background noise, as well as anisotropic distribution of noise across the sensor characteristic for any CMOS sensor.

By measuring the decaying bioluminescence signal using an EMCCD in the present experiment, after removal of the BLmini GRIN lens from the field of view, we were able to verify the areas where we should anticipate the strongest signal (Fig. 4B). The area showing highest SNR within miniscope frames aligned well with the peak bioluminescence intensity area identified via EMCCD, and corresponds to the viral injection site verified via hist logy (Fig. 4C).

## IV. Discussion

The data shown here, enabled by substantial modification of miniscope design, confirm that standard bioluminescent signals generated *in vivo* can be reliably detected. Further, the proposed benefits of design modification were realized: the BLmini is more sensitive, lighter, and simpler to assemble. Given the ongoing molecular innovation in bioluminescent indicators, and in miniscope sensitivity and resolution, this area of technology development is well-positioned to fully take advantage of the benefits of the bioluminescent strategy.

One key next step will be the refinement of spatial resolution. While being able to reliably capture temporal dynamics of bioluminescence, the design presented here offers limited spatial resolution, and can be further improved by using better imaging sensors, improved transmission optics and brighter bioluminescence constructs. CMOS sensors offering higher sensitivity are widely available and, as seen in Fig. 1C, optimization around the objective GRIN lens alone can help to further mitigate 24% of light loss. We did not attempt calcium indicator detection in the current study, focusing instead on this initial existence proof. Another key next step will be systematic testing of such signals.

Reduction in power consumption offered by removal of the excitation LED offers even greater experimental benefits for wire-free miniscope imaging. The version of the miniscope utilized for present work relied on ultrathin coaxial cable to communicate between the data acquisition system and the miniscope. However, an alternative design has recently been released, which allows wire-free collection of imaging data by replacing a wired data link with an SSD card. [9] This wire-free design is compatible with the simplified optics presented here. Further, wire-free fluorescence miniscope requires ~296 mW. Approximately half of this power is consumed by the current driver powering the LED, so its removal cuts the power consumption in half and prolongs battery life. A 45 mAh LiPo battery weighing 1.1 g allows continuous fluorescence imaging for ~20 min. When using bioluminescence indicators, this time can be extended up to 40 min, or the battery weight can be reduced to ~0.5 g to offer the same experimental duration of 20 min.

In the present report we demonstrate early results on detectability of bioluminescence using a modified UCLA miniscope optimized for bioluminescence imaging (BLmini). Removal of the excitation light, allowed us further to reduce the mass, cost, power consumption and assembly complexity of miniscopes. Even with the minimal modifications presented here, we were able to capture temporal dynamics that typ cally can be observed only when using a high-end EMCCD camera. Not only were we able to detect bioluminescence using much simpler, smaller, lower cost camera, we were able to do so at much higher framerate (as fast as 5 FPS compared to 0.1 FPS used with EMCCD [10]), paving the way towards measurements of calcium dynamics via bioluminescence indicators undergoing active development. Transitioning to bioluminescence miniature microscopy can enable testing of a broader spectrum of hypotheses.

## Supporting information

Movie of bioluminescence plotted in Figure 4AB

## Acknowledgments

Authors would like to thank Daniel Aharoni for support and valuable guidance in miniscope optimization. We also thank Kimani Toussaint for valuable discussions in optical design and testing of miniscopes, as well as Jill Juneau for discussions about electrical circuit design.

